# The Genome of the Soybean Gall Midge (*Resseliella maxima*)

**DOI:** 10.1101/2023.02.10.528044

**Authors:** Gloria Melotto, Megan W. Jones, Kathryn Bosley, Nicole Flack, Lexi E. Frank, Emily Jacobson, Evan J. Kipp, Sally Nelson, Mauricio Ramirez, Carrie Walls, Robert L. Koch, Amelia R. I. Lindsey, Christopher Faulk

## Abstract

The cecidomyiid fly, soybean gall midge, *Resseliella maxima* Gagné, is a recently discovered insect that feeds on soybean plants in the Midwest US. *Resseliella maxima* larvae feed on soybean stems which may induce plant death and can cause considerable yield losses, making it an important agricultural pest. From three pools of 50 adults each, we used long-read nanopore sequencing to assemble a *R. maxima* reference genome. The final genome assembly is 206 Mb with 64.88X coverage, consisting of 1009 contigs with an N50 size of 714 kb. The assembly is high quality with a BUSCO score of 87.8%. Genome-wide GC level is 31.60% and DNA methylation was measured at 1.07%. The *R. maxima* genome is comprised of 21.73% repetitive DNA, which is in line with other cecidomyiids. Protein prediction annotated 14,798 coding genes with 89.9% protein BUSCO score. Mitogenome analysis indicated that *R. maxima* assembly is a single circular contig of 15,301 bp and shares highest identity to the mitogenome of the Asian rice gall midge, *Orseolia oryzae* (Wood-Mason). The *R. maxima* genome has one of the highest completeness levels for a cecidomyiid and will provide a resource for research focused on the biology, genetics, and evolution of cecidomyiids, as well as plant-insect interactions in this important agricultural pest.

## Introduction

The soybean gall midge, *Resseliella maxima* Gagné (Diptera: Cecidomyiidae), is a recently discovered insect pest of soybean plants (Figure 1A) (Gagné et al. 2019). This insect was first described in 2019 after being associated in the prior year with dying soybean plants in the Midwest US (Gagné et al. 2019). Soybean plants become susceptible to *R. maxima* infestation during early vegetative growth stages, when natural fissures develop below the cotyledonary node (McMechan et al. 2021). These fissures are where the *R. maxima* females are suspected to lay their eggs (McMechan et al. 2021). After hatching, *R. maxima* larvae start feeding within the stem at the base of the plant (Figure 1B). This feeding results in necrotic lesions at the base of the plant (Figure 1C), which often results in wilting, lodging, or plant death (McMechan et al. 2021).

**Figure 1.**
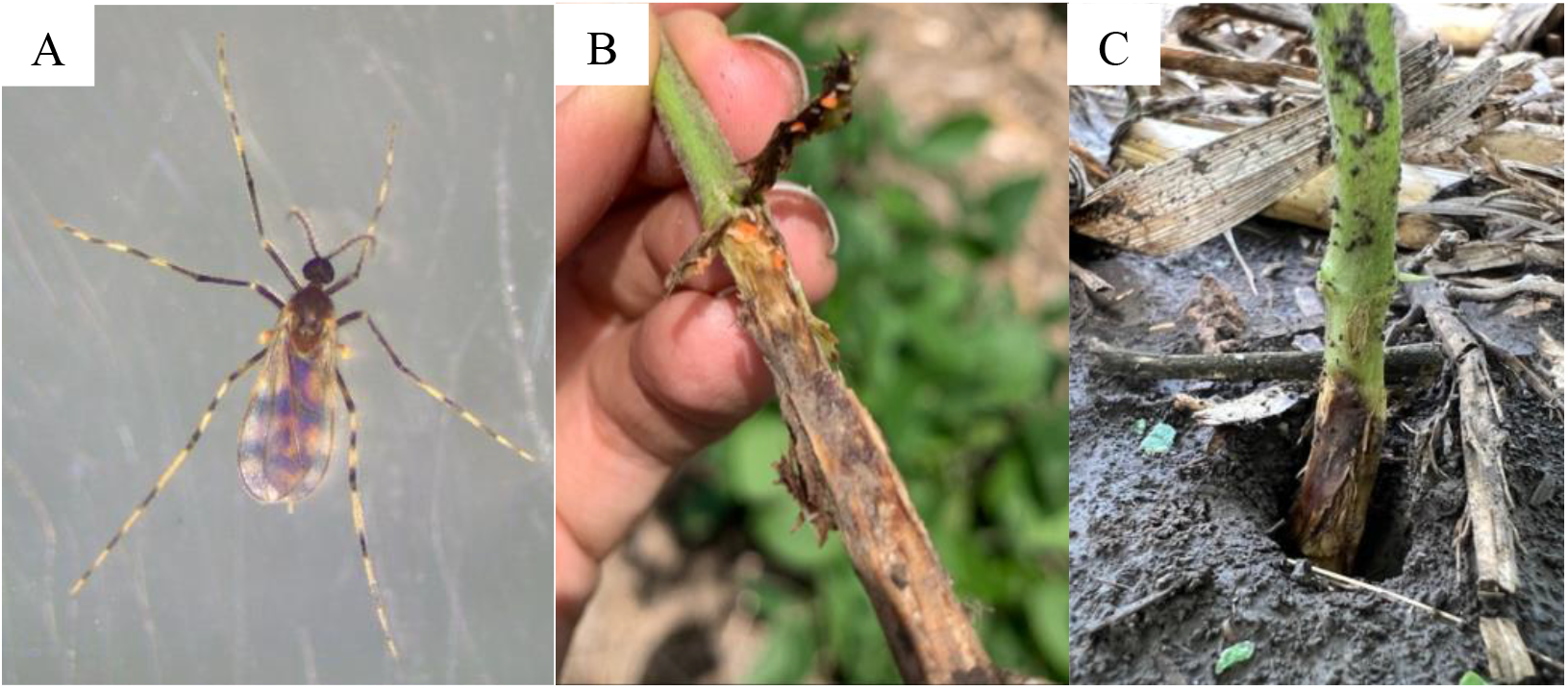
Soybean gall midge biology. **A**) Adult female of *R. maxima*. **B**) *R. maxima* larvae on soybean plant lesion. **C**) Soybean plant showing symptoms of *R. maxima* infestation in the field.

Since initial reports of *R. maxima* in 2018, the midge’s presence has been confirmed in five Midwest states: Iowa, Minnesota, Nebraska, Missouri, and South Dakota (McMechan et al. 2021). These states are ranked as the 2^nd^, 3^rd^, 4^th^, 6^th^, and 8^th^ most productive states, respectively, for soybean in the US (USDA, 2022). Soybean is a source of food and fuel, and it is an important commodity crop worldwide. In the US, soybean production accounted for 20% ($48.6 billion) of US crop cash receipts in the calendar year 2021 (USDA 2022). With *R. maxima* capable of causing yield losses (Helton et al., 2022; McMechan et al., 2021), there is growing concern over the spread and impacts of this new insect pest.

Here we provide the first genome sequence for the genus *Resseliella. Resseliella maxima* poses a threat to the US soybean industry and its genome sequence will assist with (1) evaluating biological characteristics, (e.g., overwintering ability or interactions with host plants), (2) understanding mechanisms of pesticide resistance, (3) describing cecidomyiid evolution across natural histories and host ranges, as well as (4) generating tools for accurate identification.

## Results and Discussion

### Assembly

Three pools of 50 adult individuals of *R. maxima* were digested for DNA extraction and sequenced over three flow cells on an Oxford Nanopore MinION sequencer (Supplementary File 1). A total of 13.7 Gb sequence was generated, with an N50 of 3,485 bp, and 76% of bases had a greater than Q20 quality score (Table 1). All bases were used for genome assembly. A draft assembly was generated containing 2,613 contigs with a length of 211 Mb. Decontamination and quality control filtering removed short contigs and those with anomalous coverage (<2X and >200X). The final assembly was 206 Mb, spread over 1,009 contigs with an N50 of 714,500 bp, and coverage of 64.88X. The genome-wide GC level is 31.60%. The assembly is available under NCBI Project number PRJNA928452 and accession number JAQOWM000000000

**Table 1.** Assembly statistics

| Assembly | Contigs | Length (bp) | Min length | Avg length | Max length | N50 |
| --- | --- | --- | --- | --- | --- | --- |
| Draft | 2,613 | 211,232,549 | 384 | 80,839 | 6,309,645 | 698,470 |
| Final | 1,009 | 206,036,186 | 8,001 | 204,198 | 6,309,645 | 714,500 |

| BUSCO | Complete (%) | Single (%) | Duplicate (%) | Fragment (%) | Missing (%) | n |
| --- | --- | --- | --- | --- | --- | --- |
| Draft | 87.9 | 85.1 | 2.8 | 1.9 | 10.2 | 3285 |
| Final | 87.8 | 85.2 | 2.6 | 1.9 | 10.3 | 3285 |

BUSCO scores indicated high completeness of the assembly, with no BUSCO genes lost during generation of the final assembly. The reduction of 0.1% composite score in the final assembly as compared to the draft is explained by removal of 0.2% duplicate BUSCOs and an increase of 0.1% single copy BUSCOs during assembly polishing with medaka and removal of contaminants. The final assembly has the second highest single copy BUSCO score of cecidomyiids, only exceeded by *Catotricha subobsoleta* (Alexander) (Diptera: Cecidomyiidae) (Figure 2).

**Figure 2.**
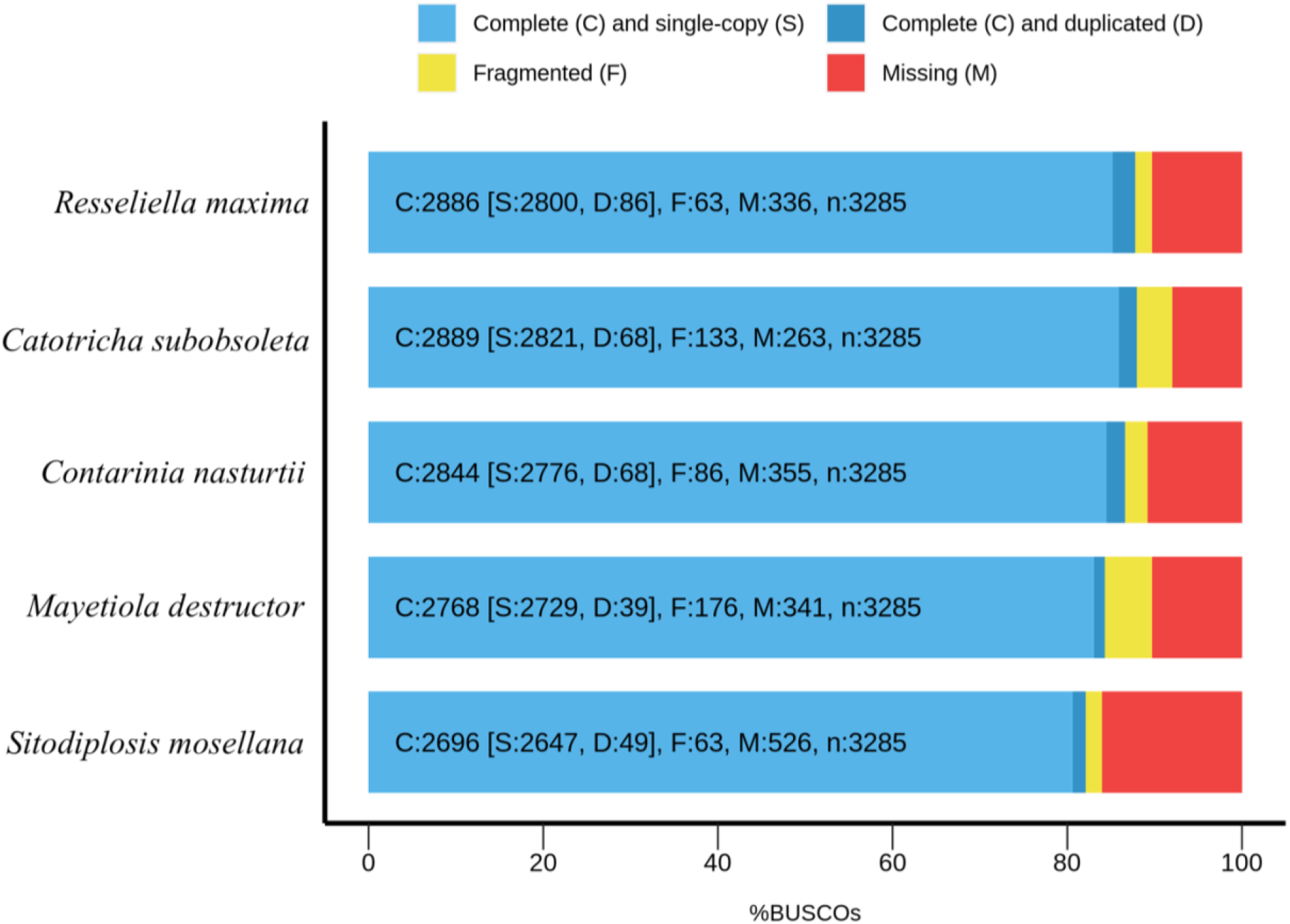
BUSCO scores for *R. maxima* assembly compared to other cecidomyiid genomes available from NCBI.

Our final assembly was 206 Mb, in line with other cecidomyiid genomes, such as *C. subobsoleta* (277 Mb), *C. nasturtii* (186 Mb), *M. destructor* (186 Mb), *P. nigripennis* (286 Mb), and *S. mosellana* (181 Mb). Assembly contiguity is high, with an N50 of 714 kb and the genome BUSCO score indicates our assembly has one of the highest completeness levels for a cecidomyiid. In addition to utility as a guide for genome assembly of related taxa, the *R. maxima* genome will contribute to a broader understanding of major biological traits associated with herbivory such as host adaptation, detoxification, and immunity (e.g., (Grbić et al., 2011), (Sparks et al., 2020), (Guan et al., 2021)).

### DNA Methylation

Nanopore sequencing can distinguish 5’methylated cytosines (5mC) from unmethylated cytosines through base calling using a methylation-aware neural network model. Global DNA methylation of cytosines in a CpG context was estimated by the fraction of 5’methylated cytosines divided by unmethylated cytosines, for all reads that aligned to the final assembly. Genome-wide methylation was 1.07%. The methylation was uniform across the assembly with only a single contig averaging above 2% (Supplementary File 2).

We saw extremely low levels of DNA methylation across the assembly, with most contigs averaging at the 1% level. While this could be a biologically meaningful level of methylation, other Diptera also have negligible levels of methylation, presumably due to loss of DNA methyltransferases (DNMT1 and DNMT3) in the dipteran common ancestor (Glastad et al., 2011). In insects with functional DNA methylation pathways, the levels are between 3-40% (Bewick et al., 2016). The level measured here could be a reflection of non-specific background DNA damage plus uncertainty in the neural network base calling model that detects DNA methylation especially at such low levels. It is unlikely that *R. maxima* contains a functional methylation pathway based on its evolutionary history.

### Repetitive DNA

To compare repeats across species, we first masked six focal cecidomyiid genomes against the most comprehensive public repeat database from the Dfam consortium. Across all cecidomyiids, repeats are poorly annotated in the existing database, as reflected by low percentages of repeat detection (Table 2). We then used RepeatModeler2, an *ab initio* repeat finding pipeline that does not rely upon prior consensus libraries which is capable of detecting repeats within a genome using only its assembly sequence. Within the *R. maxima* genome, 21.73% of the genome was masked. Other cecidomyiid genomes had repeat content ranging from 11.89% to 29.78%. Reasoning that some repeats may be more or less shared between species, we used the repeat library we generated from *R. maxima* by RepeatModeler2 to search the other cecidomyiid genomes. We found that wheat midge, *Sitodiplosis mosellana* (Géhin) (Diptera: Cecidomyiidae), shared the most repeats with *R. maxima*, followed by swede midge, *Contarinia nasturtii* Kieffer (Diptera: Cecidomyiidae), and *C. subobsoleta* and Hessian fly, *Mayetiola destructor* (Say) (Diptera: Cecidomyiidae), in that order. Despite having a high repeat content *Porricondyla nigripennis* (Meigan) (Diptera: Cecidomyiidae) shared the fewest repeats with *R. maxima* (2.88%).

**Table 2.** Repetitive content of cecidomyiid genomes

| Species | Assembly | Common name | vs.<br>Dfam<br>(%) | vs.<br>Self<br>(%) | vs.<br>SGM<br>(%) | Size<br>(Mb) |
| --- | --- | --- | --- | --- | --- | --- |
| *Resseliella maxima* | Resmax_1 | soybean gall midge | 8.15 | 21.73 | NA | 206 |
| *Contarinia nasturtii* | AAFC_CNas_1.1 | swede midge | 2.30 | 13.25 | 6.04 | 186 |
| *Porricondyla nigripennis* | ASM2654663v1 | NA | 2.26 | 18.95 | 2.88 | 285 |
| *Sitodiplosis mosellana* | ASM2101890v1 | wheat midge | 3.28 | 17.26 | 7.01 | 181 |
| *Mayetiola destructor* | Mdes_1.0 | Hessian fly | 3.47 | 11.89 | 4.46 | 186 |
| *Catotricha subobsoleta* | ASM1163474v2 | NA | 5.51 | 29.78 | 5.57 | 277 |

Overall, the genome of *R. maxima* is comprised of 15.88% interspersed repeats, 5.07% simple repeats, and 1.52% low complexity repeats (Table 3). Most interspersed repeats remain unclassified, similar to the other cecidomyiids. The full repeat complement is available in Supplementary File 3.

**Table 3.** Interspersed repeats in *R. maxima*

| Name | Number | Length<br>(bp) | Percent<br>(%) |
| --- | --- | --- | --- |
| **Retroelements** | 12207 | 8108723 | 3.94 |
| Penelope Class | 117 | 29589 | 0.01 |
| LINE Class | 8858 | 5531322 | 2.68 |
| L2/CR1/Rex | 5902 | 3795681 | 1.84 |
| RTE/Bov-B | 2250 | 1427140 | 0.69 |
| LTR Class | 3349 | 2577401 | 1.25 |
| BEL/Pao | 902 | 975078 | 0.47 |
| Ty1/Copia | 645 | 559287 | 0.27 |
| Gypsy/DIRS1 | 1802 | 1043036 | 0.51 |
| **DNA transposons** | 12991 | 5092926 | 2.47 |
| hobo-Activator | 1428 | 588862 | 0.29 |
| Tc1-IS630-Pogo | 8110 | 3313329 | 1.61 |
| MULE-MuDR | 61 | 27086 | 0.01 |
| Other | 55 | 25938 | 0.01 |
| **Rolling-circles** | 283 | 137313 | 0.07 |
| **Unclassified** | 82923 | 17722456 | 8.6 |
| **Total interspersed** |  | 30924105 | 15.01 |
| **Small RNA** | 453 | 142243 | 0.07 |
| **Simple repeats** | 252445 | 10436422 | 5.07 |
| **Low complexity** | 57017 | 3135087 | 1.52 |
| **Bases masked** |  | 44775170 | 21.73 |

Repeats in cecidomyiid genomes are poorly annotated in public libraries. Therefore, we used *ab initio* methods to identify repeats using only the genome itself and found the percentage of repeats detected increased from 8.15% to 21.73%. Reasoning that repeats are likely shared between closely related cecidomyiids, we used the *R. maxima* specific repeat library to search other genomes. Unexpectedly, we found that other Cecidomyiidae did not share a large percentage of repeats. As Cecidomyiidae is approximately 150 million years old (Dorchin et al., 2019), and very little is known about the evolutionary dynamics of this group, it is unclear if the differences in repeats are reflective of this divergence time, or, if repeats are particularly active in this group.

### Gene annotation

Putative coding regions were predicted using Gene Model Mapper (GeMoMa) against the repeat masked version of the assembly. GeMoMa uses annotations of related species as hints to detect coding regions. We used *Drosophila melanogaster* Meigan (Diptera: Drosophilidae) and *C. nasturtii* as source annotations. GeMoMa predicted 14,798 proteins with a protein BUSCO score of 89.9 %, similar to the swede midge’s score of 92.0%. The full set of proteins and associated data is available in Supplementary File 4.

Gene annotation is only available for a single cecidomyiid reference genome, *C. nasturtii*. Fortunately, that species is relatively closely related to *R. maxima* and provides good hints to the GeMoMa annotation pipeline used here. Our count of 14,798 protein coding genes is typical of an animal genome. The *C. nasturtii* assembly was created using Illumina short read sequencing with 57X coverage, while our *R. maxima* relied solely on long-read nanopore reads. Although nanopore reads are inherently less accurate than Illumina reads, we were still able to generate an accurate consensus, and its BUSCO protein score of 89.9% indicates good resolution of the assembly. Other nanopore-only assemblies have shown overall quality scores of Q45 (1 in 50,000 base error rate) (Flack et al., 2022), considered as a platinum status genome by the Vertebrate Genomes Project (Morin et al., 2020).

### Whole-Genome Phylogeny

We compared our assembly using an alignment-free whole genome-based reconstruction method yielding the tree in figure 3. We used genomes of the same six cecidomyiids as for repeat detection (plus *D. melanogaster* as an outgroup) as these are the only full genomes publicly available from Cecidomyiidae. Here *R. maxima* is sister to *S. mosellana* and *C. nasturtii* which matches the repeat content analysis where these three share the most repeats.

**Figure 3.**
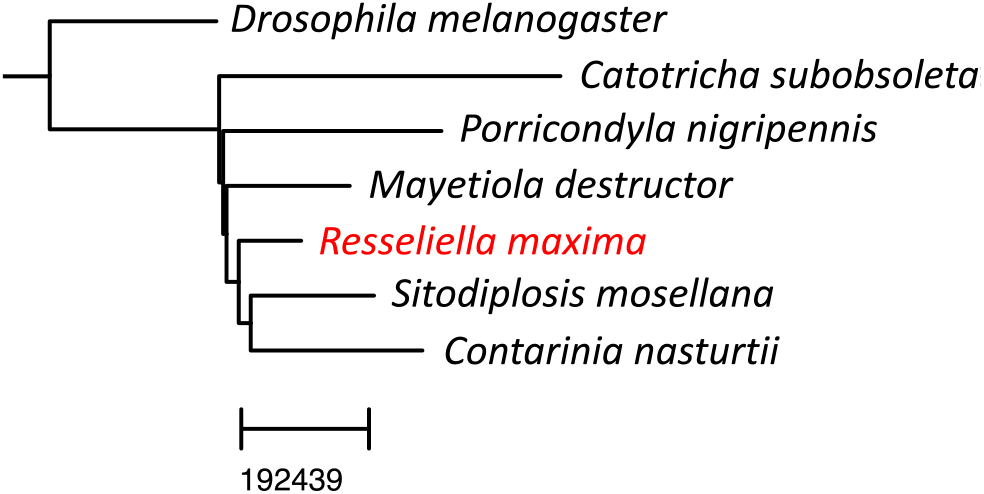
Whole genome phylogeny.

### Mitochondrial Structure and Phylogeny

We extracted the *R. maxima* mitogenome from a subset of the total reads that were identified by BLAST as homologous to the mitogenome of the Asian rice gall midge, *Orseolia oryzae* (Wood-Mason) (Diptera: Cecidomyiidae). The assembly is a single circular contig of 15,301 bp and matches 79.79 % identity over 93% of its length to the *O. oryzae* mitogenome (KM888183.1). Gene annotation indicates some errors in assembly, likely due to polymorphisms within the pooled population used for sequencing (Figure 4). The mitogenome has been deposited under accession number OQ342780.

**Figure 4.**
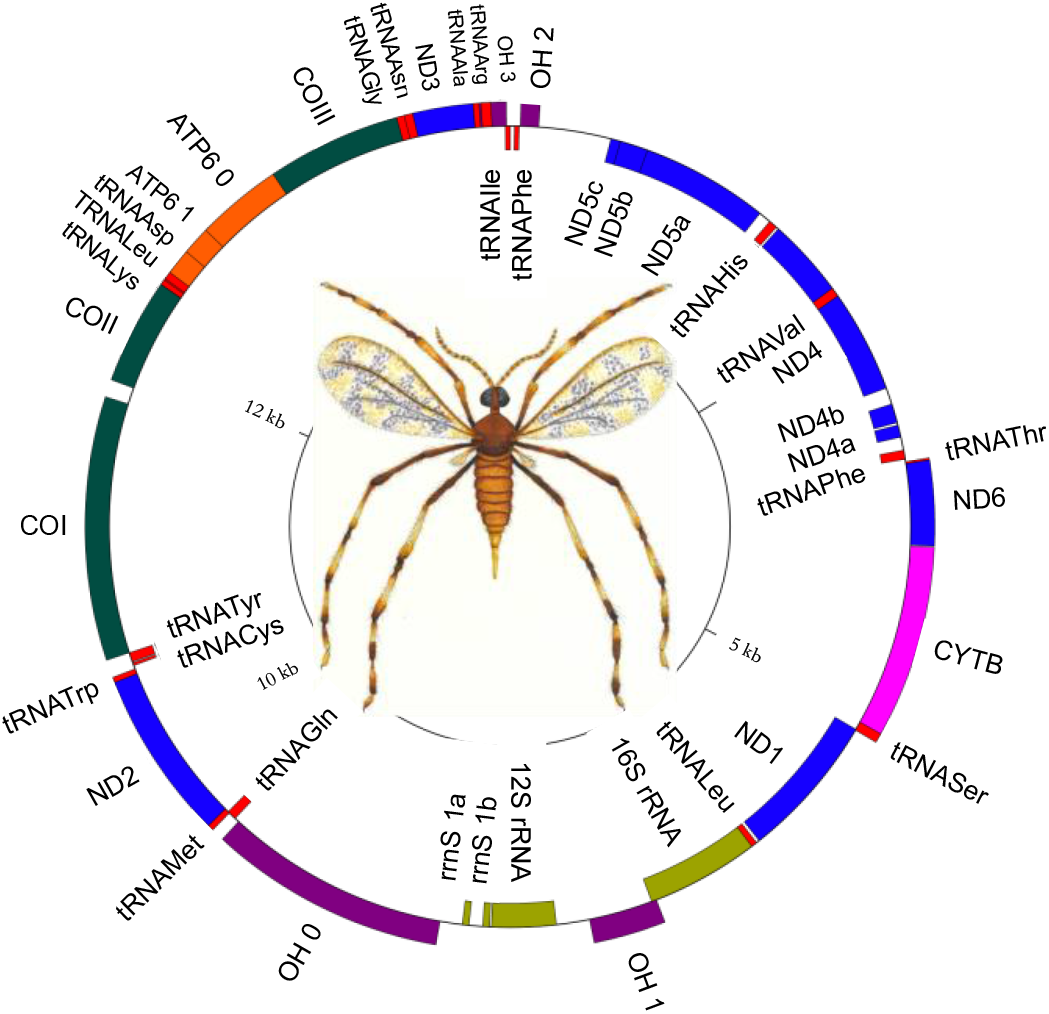
Mitochondrial structure of R. maxima.

The *R. maxima* mitogenome contains 22 tRNA genes. tRNA^Leu^ and tRNA^Ser^ have been duplicated, as in *Orseolia oryzae* and other gall midges. We were unable to annotate tRNA^Glu^ in *R. maxima*, despite multiple attempts in MITOS2. This is potentially due to the unusually truncated tRNA genes observed in other Cecidomyiid midges (Mori et al., 2021).

We compared *R. maxima* mitochondrial gene order to the closest relative for which a complete mitogenome is available, *Orseolia oryzae*. Gene order in *R. maxima* varies significantly from *O. oryzae*, with some conserved elements. Nad4 and nad5 have been inverted in both *O. oryzae* and *R. maxima* (Supplementary File 5). A region containing COII, tRNA^Lys^, atp8, atp6, and COIII is coded on the positive strand in *R. maxima*, while this entire region is inverted in *O. oryzae*. Additionally, tRNA^Leu^ and tRNA^Asp^ are present within this contiguous region in *R. maxima*, but not in *O. oryzae*. Another contiguous region on the positive strand containing tRNA^Thr^, nad6, cytB, and tRNA^Ser^ is present in both midges. With the exception of these relatively conserved regions, gene order is not well conserved between *R. maxima* and *O. oryzae*

We used 46 cytochrome oxidase I (COX1) sequences to reconstruct Cecidomyiidae relationships (Supplementary File 5.). The resulting tree placed *R. maxima* as sister to *Resseliella oleisuga* (Targioni-Tozzetti) (Diptera: Cecidomyiidae) with 88% bootstrap support. *Resseliella maxima* and *R. oleisuga* are grouped together with other *Resseliella* species in a polyphyletic clade that includes other gall midges in the subtribe Cecidomyiini, as well as gall midges in Aphidoletini, Lopesiini, and Lestodiplosini. These polyphyletic clades are likely a result of (1) a single-locus phylogenetic reconstruction, (2) the significant changes we see in mitochondrial genomes across Cecidomyiidae, and/or (3) the poor coverage of available cecidomyiid mitogenomes. However, this could potentially indicate a need for reexamination of cecidomyiid phylogeny.

Phylogenies were performed in two different ways due to the availability of comparative data. First, we created a phylogeny based on whole genome comparisons and found *R. maxima* most closely related to *S. mosellana and C. nasturtii*. In contrast, Dorchin et al.’s multi-locus phylogeny places at least one other Resseliella species more closely to *C. nasturtii* before *S. mosellana*. Importantly, our phylogeny was limited to seven species for which whole genomes exist. Secondly, we created a larger phylogeny using a database of COX1 amino acid sequences and found that *R. maxima* grouped with several other *Resseliella* but that the genus appears to be polyphyletic overall. Our mitogenome likely contains a few errors as assembly was particularly difficult in light of pooled sampling, however, others have shown nanopore-only mitogenomes are useful and reliable despite imperfections (De Vivo et al., 2022).

### Absence of Wolbachia infection

Some members of Cecidomyiidae host *Wolbachia* as intracellular bacteria (Behura et al., 2001; Bel Mokhtar et al., 2020). To test whether *R. maxima* were infected, we examined the sequencing reads for the presence of *Wolbachia*. Minimap2 was used to identify putative candidates in the total set of sequenced reads by matching to the full genomes of a set of nine strains that cover a wide range of *Wolbachia* diversity. There were 4929 sequences (0.000036%) that matched these reference *Wolbachia* genomes. Of these 92% (n=4,480) were identified as human origin, and the remaining 29 hits matched to a variety of non-*Wolbachia* bacteria (Supplementary File 6). This evidence indicates that this population of *R. maxima* is not hosting *Wolbachia* infection. As validation, we performed PCR on genomic isolates and found no amplification using general *Wolbachia* primers.

### Summary

Here we provide the first whole-genome assembly of a *Resseliella* species (206 Mb with 64.88X coverage) that has one of the highest completeness levels (BUSCO score of 87.8%) for a cecidomyiid. The *R. maxima* genome will provide a resource for research focused on the biology, genetics, and evolution of cecidomyiids, as well as development of management tactics.

Genomes of insects and other invertebrates can be difficult to assemble for a range of factors including: (1) high heterozygosity when inbreeding is unrealistic, (2) small sizes which present very little DNA, requiring pooling individuals to make libraries, (3) lack of high quality genome assemblies of close relatives to assist with assembly due to high diversity across arthropods, (4) the need to optimize DNA preparations for new insect species, and (5) the fact that some arthropods have large genomes (Richards and Murali, 2015). However, long-read sequencing technologies (e.g., Pacific Biosciences (PacBio) and Oxford Nanopore) are contributing to improvement in quality of genome assembly, producing assemblies ~48X more contiguous than short-read based approaches (Hotaling et al., 2021; Li et al., 2019). Other pest insect genomes have recently been sequenced using nanopore-only approaches such as the black carpenter ant (*Camponotus pennsylvanicus* (De Geer), Hymenoptera: Formicidae) and the coconut rhinoceros beetle (*Oryctes rhinoceros* [L.] (Coleoptera: Scarabaeidae)) (Faulk, 2022; Filipović et al., 2022). Here we not only assembled a high-quality genome from long reads, but were able to do so from pooled samples.

Knowledge of arthropod pest genomes can reveal important evolutionary innovations. For example, the genome of the spider mite, *Tetranychus urticae* Koch (Trombidiformes: Tetranychidae), showed the evolutionary innovation of silk production, and signatures of polyphagy and detoxification (Grbić et al. 2011). Guan et al. (2021) used whole genome sequencing to detect insecticide resistance mutations in fall armyworm, *Spodoptera frugiperda* (JE Smith) (Lepidoptera: Noctuidae). Genomic analysis of the brown marmorated stink bug, *Halyomorpha halys* (Stål) (Hemiptera: Pentatomidae), revealed genetic elements associated with immunity and detoxification that have potential for biomolecular pesticide applications (Sparks et al. 2020).

Even though the family Cecidomyiidae has more than 6,600 described species (Dorchin et al., 2019) there are only five genome assemblies from this family and none from the genus *Resseliella* available on GenBank. Not surprisingly, most of the genomes belong to agricultural insect pests (i.e., *M. destructor, C. nasturtii*, and *S. mosellana*). Availability of the *R. maxima* genome will facilitate population genetics-based understandings of the origin of *R. maxima* and its spread to new areas, as well as provide possibilities for future work on developing alternative pest control methods.

## Methods

### Sample collection

*Resseliella maxima* adults were reared from field-collected soybean stems symptomatic of infestation with *R. maxima*. The collection of the stems occurred in summer of 2022 at one farm in Rock County, Minnesota, US. Symptomatic soybean plants were collected by pulling the entire plants from the soil. The plants were then trimmed above the first pair of unifoliate leaves and the roots to a length of five centimeters. Each stem was wrapped with a small piece of PARAFILM® at the cut end to decelerate plant dehydration. Ten trimmed stems were set vertically into a 3-cm deep layer of potting soil (BM2 Seed Germination and Propagation Mix, Berger, Saint-Modeste, Quebec, CA) in one emergence cage. Emergence cages consisted of plastic 5-liter clear paint mixing buckets with lids (TCP Global Corporation, Lakeside, California, USA), with a 6-cm diameter hole that had a fine-mesh (Quest Outfitters, Sarasota, Florida, USA) sleeve 30-cm long attached to it to facilitate access to the contents of the cages. The emergence cages were maintained at room temperature in 16:8 (light:dark) and watered as needed. Adult insects were collected manually into microcentrifuge tubes, freeze-killed, and morphologically confirmed to be *R. maxima* according to Gagné et al. (2019).

### DNA extraction and sequencing

DNA was extracted using a Zymo Quick-DNA Miniprep Plus kit, (catalog number D4068, Zymo Research, Irvine, CA), according to manufacturer’s instructions. Due to sample timing availability and flow cell upgrade paths, we performed sequencing with two different flow cell types over three flow cells. For each of three pools, approximately 50 individuals were used for extraction, generating 1 μg of DNA which was loaded into the library prep kit. Libraries were prepared using the SQK-LSK110 and SQK-LSK114 ligation sequencing kits for flow cells R9.4.1 and R10.4.1 respectively. Sequencing was carried out for 24 hours per flow cell. Bases were called using the guppy basecaller v6.3.9 with model ‘dna_r10.4.1_e8.2_400bps_modbases_5mc_cg_sup.cfg’.

### Genome Assembly and Polishing

*De novo* assembly of the *R. maxima* nuclear genome was accomplished using Flye v2.9 (https://github.com/fenderglass/Flye) with a subsequent polishing step done using Medaka v1.6.0 (https://github.com/nanoporetech/medaka). We used BUSCO (v5.4.3) to assess genome completeness for the draft assembly both before and after Medaka polishing steps (Manni et al., 2021). Specifically, the Diptera OrthoDB v10 database, which consists of 3,285 single-copy orthologs, was chosen for scoring our assemblies. Based on these assessments, we then selected the polished assembly with the highest BUSCO score for decontamination and downstream analyses. Full commands for all steps in the bioinformatic pipeline are given in Supplementary File 7.

### Decontamination

The blobtoolkit (https://blobtoolkit.genomehubs.org/blobtools2/) was used to examine contig properties by comparing GC content, contig length, coverage, and BLAST matches to the NCBI non-redundant (nr) database (Challis et al., 2020). When deciding cutoff values, the presence of BUSCO genes within a contig was used to determine thresholds. For instance, all contigs below 8,000 bp were removed as none below that size contained any BUSCOs. We removed contigs below 20X and above 200X coverage.

### Methylation

5’ DNA methylation at cytosines in a CpG context was assessed during initial base calling by using a DNA modification aware model. Output files were converted to bed format using modbam2bed v0.6.2 (https://github.com/epi2me-labs/modbam2bed). Aggregation of DNA methylation was performed with ‘awk’ on the command line.

### Repeats

RepeatMasker v4.1.4 (https://www.repeatmasker.org/cgi-bin/WEBRepeatMasker) with the full Dfam library v3.6 (https://www.dfam.org/home) was initially used for all cecidomyiid repeat assessments (Flynn et al., 2020; Storer et al., 2021). For ab initio repeat detection, RepeatModeler2 v2.0.2a (https://www.repeatmasker.org/RepeatModeler/) was used on each genome independently (Flynn et al., 2020). To determine shared repeat content, the *R. maxima*-specific repeat library generated from RepeatModeler2 was used as input for RepeatMasker to mask other cecidomyiid genomes.

### Protein annotation

GeMoMa v1.9 (http://www.jstacs.de/index.php/GeMoMa) was used for homology-based protein model prediction with both the *Drosophila. melanogaster* (GCA_000001215.4) and *Contarinia. nasturtii* (GCF_009176525.2) transcriptomes as references (Keilwagen et al., 2019). BUSCO was run against the resulting GeMoMa annotation in protein mode with the Diptera odb10 database to assess quality.

### Mitochondria assembly

Total reads were first blasted by mtblaster (https://github.com/nidafra92/squirrel-project/blob/master/mtblaster.py) using the *Orseolia oryzae* mitogenome (KM888183) to select for reads with high identity to a cecidomyiid mitochondria sequence. Next, resulting reads were filtered by nanofilt (https://github.com/wdecoster/nanofilt) to keep only reads above 15 kbp in length and average Q-score above 30. Finally, flye was used to perform mitogenome assembly (Kolmogorov *et al*. 2020). A single circular contig was recovered with 785X coverage. The assembly was polished using four rounds of racon polishing (https://github.com/lbcb-sci/racon) followed by medaka with the same parameters as the nuclear assembly.

### Wolbachia detection

*Wolbachia* infection was determined using the references of nine *Wolbachia* genomes broadly covering the *Wolbachia* phylogeny. The strains are described in table 4. All strain genomes were concatenated and used as the query against the set of total sequencing reads. Minimap2 was used to find high identity hits. These hits were used as input to kraken2 for species identification using the K2-Standard-16Gb database (https://benlangmead.github.io/aws-indexes/k2) version 2022-06-07 (Wood et al., 2019). Secondly, a PCR based approach was used to validate the absence of *Wolbachia*. We used *Wolbachia-specific* W-Spec primers ((Werren and Windsor, 2000) (W-Specf (CATACCTATTCGAAGGGATAG), W-Specr (AGCTTCGAGTGAAACCAATTC) to screen for the presence of *Wolbachia* 16S DNA in the sample. The PCR method failed to detect *Wolbachia*, affirming our computational findings.

**Table 4.**
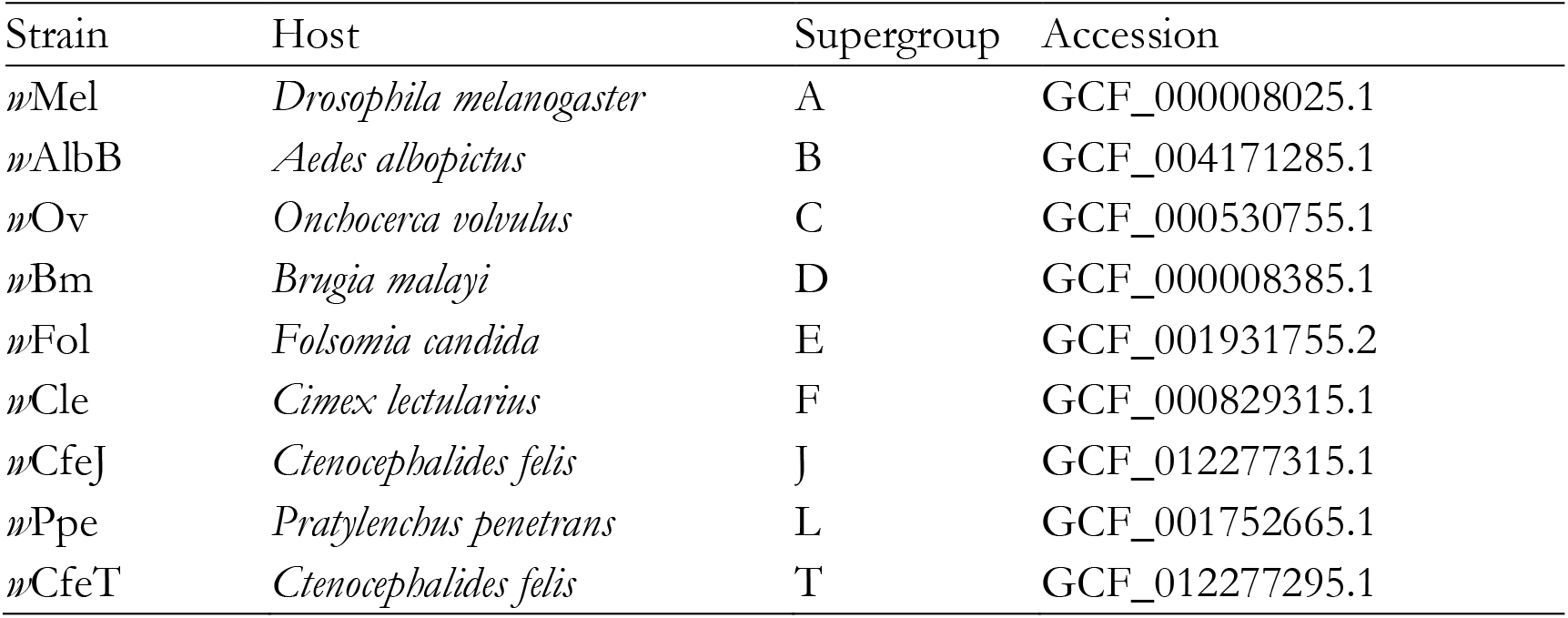
*Wolbachia* strains used for infection detection.

### Phylogenetic reconstruction

A whole genome phylogeny was created with an alignment-free method, SANS, that follows a pangenomic approach to efficiently calculate a set of splits in a phylogenetic tree or network (Rempel and Wittler, 2021). Default settings were used. A mitochondrial phylogeny was created using COX1 sequences from (Dorchin et al., 2019). Amino acid sequences were aligned using MEGA11. Since many of the COX1 sequences available on NCBI are partial and vary in length, we trimmed the aligned sequences to roughly equal size in MEGA11 (Supplementary File 5). After aligning and trimming, we assembled a phylogeny in IQ-TREE 1.6.12 with the following settings: Alignment: sgm_phylo_protein_align.fas, # of sequences = 47, Sequence type & substitution model: Amino acids, mtART, Rate heterogeneity: None, State frequency: Estimated by Maximum likelihood, Bootstrap brach support: UltraFast, # of replicates = 1000, Single branch test: None, Tree search: Perturbation strength = 0.5, # of unsuccessful iterations to stop = 500, Root tree: Outgroups: *Rabdophaga heterobia*. The tree was rooted on *Rabdophaga heterobia* (Loew) (Diptera: Cecidomyiidae), which is in the tribe Lasiopteridi. The remaining cecidomyiid COX1 sequences were from the tribe Cecidomyiidi, which includes *Resseliella* and close relatives (Dorchin et al., 2019). The mitogenome was visualized using GenomeVx on the web (Conant and Wolfe, 2008).

## Data Availability

Assembly and BioSample information is available at NCBI Project number PRJNA928452 and accession number JAQOWM000000000. The mitochondrial assembly is available separately at accession number OQ342780. Supplementary data is available at FigShare.

## Supplemental Data

**Supplementary File 1.**
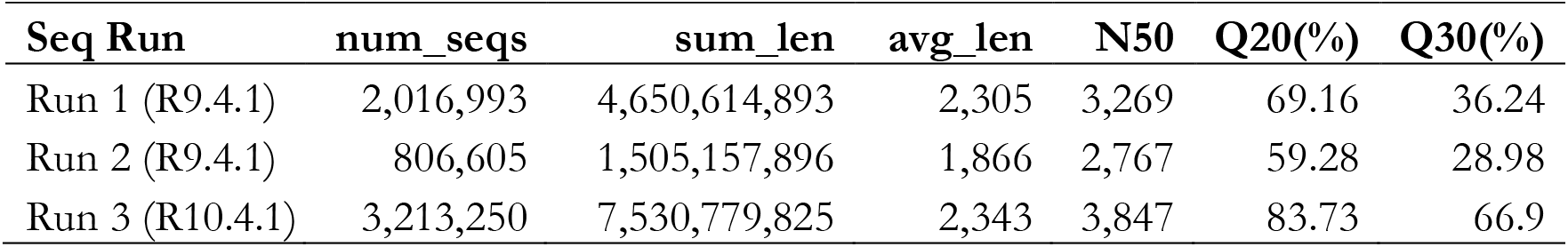
Sequencing run statistics table

| Seq Run | num_seqs | sum_len | avg_len | N50 | Q20(%) | Q30(%) |
| --- | --- | --- | --- | --- | --- | --- |
| Run 1 (R9.4.1) | 2,016,993 | 4,650,614,893 | 2,305 | 3,269 | 69.16 | 36.24 |
| Run 2 (R9.4.1) | 806,605 | 1,505,157,896 | 1,866 | 2,767 | 59.28 | 28.98 |
| Run 3 (R10.4.1) | 3,213,250 | 7,530,779,825 | 2,343 | 3,847 | 83.73 | 66.9 |

**Supplementary File 2.**

Genome-wide methylation figures

**Supplementary File 3.**

Repeat information

**Supplementary File 4.**

Protein annotation data

**Supplementary File 5.**

Mitogenome gene order comparison between *R. maxima* and *O. oryzae*. Maximum likelihood phylogenetic tree, accession numbers, trimmed alignments, and newick file of Cecidomyiidae cytochrome oxidase I (COX1) sequences from NCBI.

**Supplementary File 6.**

Kraken2 output identifying all read hits to *Wolbachia* species.

**Supplementary File 7.**

Detailed computational methods

## Author contributions

This work was a collaborative effort by the members of graduate course ANSC 8509 taught by Dr. Faulk in the Department of Animal Science at the University of Minnesota in Fall 2022. Students were Melotto, Jones, Bosley, Flack, Frank, Jacobson, Kipp, Nelson, Ramirez, and Walls. Students performed analyses and provided text for the manuscript. Drs. Lindsey and Koch provided the midge samples, analytical guidance, and edited the manuscript. Dr. Faulk performed analyses and edited the manuscript. All authors have read and approved the manuscript prior to submission.

